# Genetic-based inference of densities, effective and census sizes of expanding riverine meta-populations of an invasive large-bodied freshwater fish (*Silurus glanis L.*)

**DOI:** 10.1101/2024.04.05.588309

**Authors:** Ivan Paz-Vinas, Géraldine Loot, Stéphanie Boulêtreau, Marlène Chiarello, Charlotte Veyssière, Jessica Ferriol, Frédéric Santoul

## Abstract

Effective (N_e_) and census (N_c_) population sizes are key eco-evolutionary parameters. Jointly estimating them have an important practical value for efficient conservation and wildlife monitoring and management. Assessing N_e_ and N_c_ remains however challenging for elusive, rare species or species inhabiting in complex habitats like large rivers. Genetic-based N_e_ estimations could help resolve complex situations, as only a handful of genotyped individuals are needed to estimate N_e_, and then N_C_ can be subsequently using an N_e_/N_C_ ratio. However, most N_e_ estimation methods are based on restrictive assumptions (e.g. Wright-Fisher model) making them inappropriate for inferring N_e_ and N_c_ for populations and species exhibiting complex dynamics. Here, we aimed at estimating N_e_, N_C_ and densities for meta-populations of a large invasive freshwater fish (the European catfish *Silurus glanis*) that has been introduced in the Garonne-Dordogne river basin (Southwestern France), using a framework that combines multiple data sources and approaches. First, we characterized spatial patterns of genetic variation using microsatellite genotype data, revealing a significant isolation by distance pattern informing about the species’ dispersal capacities. We then detected four genetically-distinct clusters of individuals coexisting in the river basin that might be the result of multiple introductions from different genetic sources. Further, we characterized the demographic expansion of the species at the river basin scale by analyzing data from a multidecadal demographic monitoring survey, and estimated a specific Ne/Nc ratio for this species. We finally combined all the gathered information to design four competing demo-genetic models accounting for all the complexity of *S. glanis* meta-populations inhabiting the river basin. We simulated data under these models and then inferred Ne, Nc and densities through approximate Bayesian computation and random forest procedures. We show how multiple genetic and non-genetic approaches can be combined to estimate N_e_ and N_c_ in hard-to-monitor meta-populations exhibiting complex demo-evolutionary dynamics.

## INTRODUCTION

Estimating effective (N_e_) and census (N_c_) population sizes is of prime importance for efficient wildlife management (Schwartz et al., 1998; Luikart et al., 2010; Palstra and Fraser, 2012; Pauls et al., 2014; Ryman et al., 2019; Hoban et al., 2020; Clarke et al., 2023). N_e_ is a key genetic parameter that informs about the long- and short-term viability of populations (Garner et al., 2020; Hoban et al., 2020; Laikre et al., 2020; Hoban et al., 2021; Waples, 2023). There is a general agreement that populations with N_e_<500 will experience increasing genetic depletion and reducing evolutionary potential with time, while populations with N_e_<50 are considered at immediate risk of extinction due to inbreeding depression and pervasive effects of genetic drift (Franklin, 1980; Soulé, 1980; Frankham et al., 2002; Jamieson and Allendorf, 2012; but see Frankham et al., 2014 and Clarke et al., 2023 for discussion on revised thresholds recommendations). Effective population size has also recently entered the global conservation policy arena, given that the N_e_ 500 threshold is now at the core of one of the 26 headline indicators of the Monitoring Framework of the UN Convention on Biological Diversity (CBD) Kunming-Montreal Global Biodiversity Framework (i.e. headline indicator A.4; CBD, 2022; Hoban et al., 2024). Concomitantly, N_c_ may indicate at which level populations are exposed to demographic and environmental stochastic events that may lead to drastic changes in recruitment patterns or population sizes (Palstra and Fraser, 2012). Assessing both N_e_ and N_c_ is thus particularly useful from a biodiversity management standpoint, as these indicators allow monitoring genetic and demographic changes in populations, evaluating their viability at different timescales, identifying priority populations for designing appropriate conservation actions and informing policy (Luikart et al., 2010; Garner et al., 2020; Laikre et al., 2020; Hoban et al., 2020; Laikre et al., 2021; Frankham, 2021).

Although being generally conducted from a conservation standpoint (e.g. to identify at-risk populations), joint N_e_ and N_c_ assessments may also have an important practical value in the context of biological invasions monitoring and management, as they may help identify, monitor and prioritize invasive populations requiring active population control actions at local scales (Blanchet, 2012; Paz-Vinas et al., 2021). For instance, introduced populations with N_e_<50 can be expected to be less likely to establish, be maintained and become invasive compared to introduced populations with N_e_>500, due to the expected pervasive genetic effects associated with low effective population sizes and genetic diversities (e.g. increased inbreeding depression or accumulation of deleterious alleles; Frankham et al., 2002; Jamieson and Allendorf, 2012). Depending on the socioeconomic and ecological contexts, biological invasion control measures may target populations (i) with high N_e_, e.g. by increasing intentionally their extinction risk through removal campaigns aimed at reducing their N_e_ (when total eradication is unfeasible), or (ii) with low N_e_, because they are theoretically more prone to become extinct, and hence more prone to successfully respond to population control measures than populations with high N_e_. However, many invasive populations succeed despite displaying very low levels of genetic diversity and N_e_ values (i.e. the “genetic paradox” dilemma in invasion biology; Roman and Darling, 2007; Rollins et al., 2013). The adoption of efficient management actions must therefore also take into account N_c_, as invasive populations displaying a large N_c_ are likely to produce large amounts of propagules that might increase their invasion success (Lombaert et al., 2010; Bertelsmeier and Keller, 2018; Javal et al., 2019).

Assessing N_e_ and N_c_ remains however challenging for highly mobile, cryptic, nocturnal, shy and/or rare species for which direct estimation protocols (e.g. based on direct counts) are inefficient or inapplicable (Luikart et al., 2010; Laikre et al., 2021). This is the case for many aquatic, forest-dwelling, or subterranean species for which active and passive counting cannot be implemented due to unfavourable environmental conditions and/or to particular life-history traits affecting species’ observability, catchability, or occurrence frequencies. For instance, direct counting in rivers (e.g. through snorkeling surveys or using underwater video tracking devices; Thurow and Schill, 1996; Boussarie et al., 2016) is sometimes impracticable. In such ecosystems, the relative abundance and density of fish species can be estimated by coupling different sampling methods such as electric fishing and angling with Capture-Mark-Recapture (hereafter, CMR) procedures (e.g. Mycko et al., 2018). Although very efficient, these approaches are associated with technical constraints, as they generally imply a significant sampling effort (e.g. repeated capture sessions, a high minimal number of marked individuals). Further, the efficiency of each sampling technique depends on multiple factors including species’ biological traits like fish body size (Mycko et al., 2018), and environmental parameters (e.g. water conductivity or stream depth; Allard et al., 2014). Specifically, electrofishing is inefficient in large watercourses with moderate-to-high water depths (Allard et al., 2014; > 1 meter; Murphy and Willis, 1996) and for large-bodied fish, hence preventing reliable abundance estimations in such conditions.

In these cases, population genetics provides useful tools for indirectly estimating both N_e_ and N_c_ requiring relatively small numbers of individuals (Luikart et al., 2010; Pauls et al., 2014; Hoban et al., 2020). The neutral genetic diversity of a population is indeed positively correlated to N_e_ (Nei, 1987), which is in turn related to the census size N_c_ (Hamilton, 2009). Thus, deriving N_c_ from genetic data is theoretically feasible (Nei, 1987; Frankham, 1996), provided that the relationship between N_e_ and N_c_ (i.e. the N_e_/N_c_ ratio; Palstra and Fraser, 2012; Gomez-Uchida et al., 2013) is properly known (see Hoban et al., 2020 and Frankham et al. (2019) for recent reviews of N_e_/N_c_ values across multiple species). This ratio is a key genetic parameter varying among taxonomic groups, species and populations due to differences in evolutionary trajectories, life-history traits, metapopulation structure and demographic histories (Frankham, 1995; Palstra and Fraser, 2012; Hoban et al., 2020, 2021). A consensual N_e_/N_c_ = 0.1 value is typically assumed for most taxonomic groups in the absence of knowledge of a specific ratio for a given species or taxonomic group (Hoban et al., 2020; Frankham, 2021; Laikre et al., 2021; Hoban et al., 2021, 2023, 2024).

Several methods have been developed for inferring N_e_ from genetic data (Storz and Beaumont, 2002; Waples, 2006; Waples and Do, 2008; Tallmon et al., 2008; Antao et al., 2011; Do et al., 2014; Wang et al., 2016; Santiago et al., 2020, 2024). However, most methods undertake restrictive assumptions that are not always met by the studied populations (Paz-Vinas et al., 2013; Hössjer et al., 2016; Ryman et al., 2019). Among others, most methods implicitly or explicitly assume that the populations under study meet assumptions of the Wright-Fisher model (Fisher, 1922; Wright, 1931), i.e. that the population is closed (i.e. does not exchange migrants with other populations), unstructured, and has not experienced demographic changes over time (constant population size). Although some populations may approximate this theoretical model, such highly restrictive assumptions are rarely verified in the wild. Thus, N_e_ estimation methods relying on a Wright-Fisher population model, or assuming any other equilibrium state (e.g. population being at the drift/migration equilibrium) can be biased by factors like population structure, admixture, demographic history or specific gene flow patterns (Chikhi et al., 2010; Heller et al., 2013; Paz-Vinas et al., 2013; Ryman et al., 2019). Therefore, these methods might provide limited output for guiding the management of populations experiencing complex demographic histories or displaying complex patterns of population structure like invasive populations (Ryman et al., 2019). Further, some classical N_e_ estimation approaches (e.g. linkage disequilibrium-based approaches) sometimes produce negative or infinite estimates (e.g. Paz-Vinas et al., 2013; Castagné et al., 2023), a problem that might arise due to sampling issues (Waples and Do, 2008). This impasse might preclude point derivations of N_c_ from N_e_ *via* a N_e_/N_c_ relationship.

Model-based inference procedures like approximate Bayesian computations (ABC; Beaumont et al., 2002) constitute a useful and flexible alternative approach to classical linkage-disequilibrium-based N_e_ estimation methods (Tallmon et al., 2008; Hong et al., 2023). When coupled with powerful simulation tools allowing to set up user-defined demographic and evolutionary models, ABC techniques allow the estimation of contemporary N_e_ for populations displaying complex meta-population structures (e.g. populations living in dendritic river systems; Paz-Vinas et al., 2015) and/or complex demographic histories (e.g. multiple introduction events and population size changes; Rey et al., 2015).

Here, we used neutral microsatellite genotypic data and model-based ABC-random forest procedures (ABC-RF; Pudlo et al., 2016) to estimate effective population sizes (N_e_), abundances (N_c_) and densities of meta-populations from an invasive freshwater fish species (the European catfish *Silurus glanis L.*) inhabiting four large rivers belonging to the Garonne-Dordogne hydrographical basin (Southwestern France). We first described the spatial patterns of genetic variation observed for this species across the basin. Second, we simulated genetic data under four competing population genetics models calibrated to recreate the connectivity among rivers, the hydrographical basin colonization process, the demographic changes (i.e. ongoing demographic expansion) experienced by *S. glanis* populations since its introduction in the Garonne-Dordogne basin, and the observed spatial genetic structure of *S. glanis* meta-populations. We used ABC-RF model-choice procedures to determine the model that best explains the observed patterns of genetic variation. We then inferred current N_e_ values (i) for each sampled site (i.e. within*-*site N_e_ values) and (ii) for the meta-populations identified in the four rivers (meta-N_e_ values, i.e. the effective size of each meta-population) using simulations generated under the best-supported population genetics model. Next, we compared estimates of abundance and density of European catfish individuals in our study area derived from meta-N_e_ values by using (i) a specific N_e_/N_c_ ratio estimated using repurposed CMR data and a subset of our genetic dataset, and (ii) the consensual N_e_/N_c_ = 0.1 ratio.

Our study provides an example of how a powerful analytical framework such as ABC-RF, a snapshot population genetic survey, and the analysis of demographic monitoring data can be combined to jointly infer both N_e_ and N_c_ in hard-to-monitor meta-populations exhibiting complex demo-evolutionary dynamics. The results of this study will provide key information to regional stakeholders and biodiversity managers to quantify the potential impacts of *S. glanis* on native and protected ichthyofauna predated by *S. glanis*, and to guide the design of future *S. glanis* population control measures.

## MATERIALS & METHODS

### Data collection

*Biological model.* The European catfish (*Silurus glanis L.*) is one of the twenty largest freshwater fish species worldwide (Stone, 2007; Copp et al., 2009), and the largest in Europe (> 2.7 meters and > 130 kg of weight for the biggest individuals; Boulêtreau and Santoul, 2016; Cucherousset et al., 2018). Native from Central and Eastern Europe, this species has been successfully introduced in Southern and Western Europe for recreational angling during the twentieth century (Castagné et al., 2023). In France, evidence from multiple introductions involving different genetic sources has been previously identified (Castagné et al., 2023), with successful introduction events due to deliberate releases having occurred since 1983 in the Garonne-Dordogne River basin (Castagné et al., 2023). This previous study, conducted at a European scale, suggests that multiple introductions involving multiple sources might have specifically occurred in the Garonne-Dordogne River basin, though the exact number of introductions, the location of the introduction events and the origin of the released individuals remains partially unknown. There is evidence that this opportunistic species has significant ecological impacts in the Garonne-Dordogne basin due to predation on threatened migratory species such as the Atlantic salmon *Salmo salar* (Boulêtreau et al., 2018), the sea lamprey *Petromyzon marinus* (Boulêtreau et al., 2020a), and shads *Alosa alosa* and *Alosa fallax* (Boulêtreau et al., 2021; Guillerault et al., 2017). The species has a long lifespan (>40 years), high fecundity and high behavioural plasticity, three characteristics that should have facilitated its rapid demographic expansion in the Garonne hydrographical basin (Cucherousset et al., 2018; Proteau et al., 2008).

*Sampling design.* Our study focused on the four main rivers of the Garonne-Dordogne hydrographical basin (>100,000 km^2^, Southwestern France): the Garonne, Dordogne, Lot and Tarn rivers (Figure 1). Between Spring 2015 and Summer 2018, a collaborative sampling was performed by different contributors, including our research group, local practitioners, and recreational and professional anglers, each contributor using different sampling techniques (electrofishing, fyke net traps and angling; see Table 1 for details). The use of different sampling techniques was necessary to sample individuals representative of the wide body size spectrum displayed by this species in the study area (from <10 cm to 2.73 meters; Boulêtreau and Santoul, 2016). A total of 559 individuals were sampled at 18 different sampling sites (Figure 1, Table 1). We aimed at collecting as many individuals as possible *per* site for subsequent genetic analyses, although there was high inter-site variability in the number of successfully sampled individuals (i.e. sample sizes ranged from 9 to 96 individuals; see Table 1). For each individual, a small fragment of the pelvic fin was collected and stored in 90% ethanol for subsequent laboratory analyses. All individuals were released alive at their sampling location except those captured by professional anglers (Table 1), who subsequently capitalized on sampled individuals as a commercial source of food.

**FIGURE 1:**
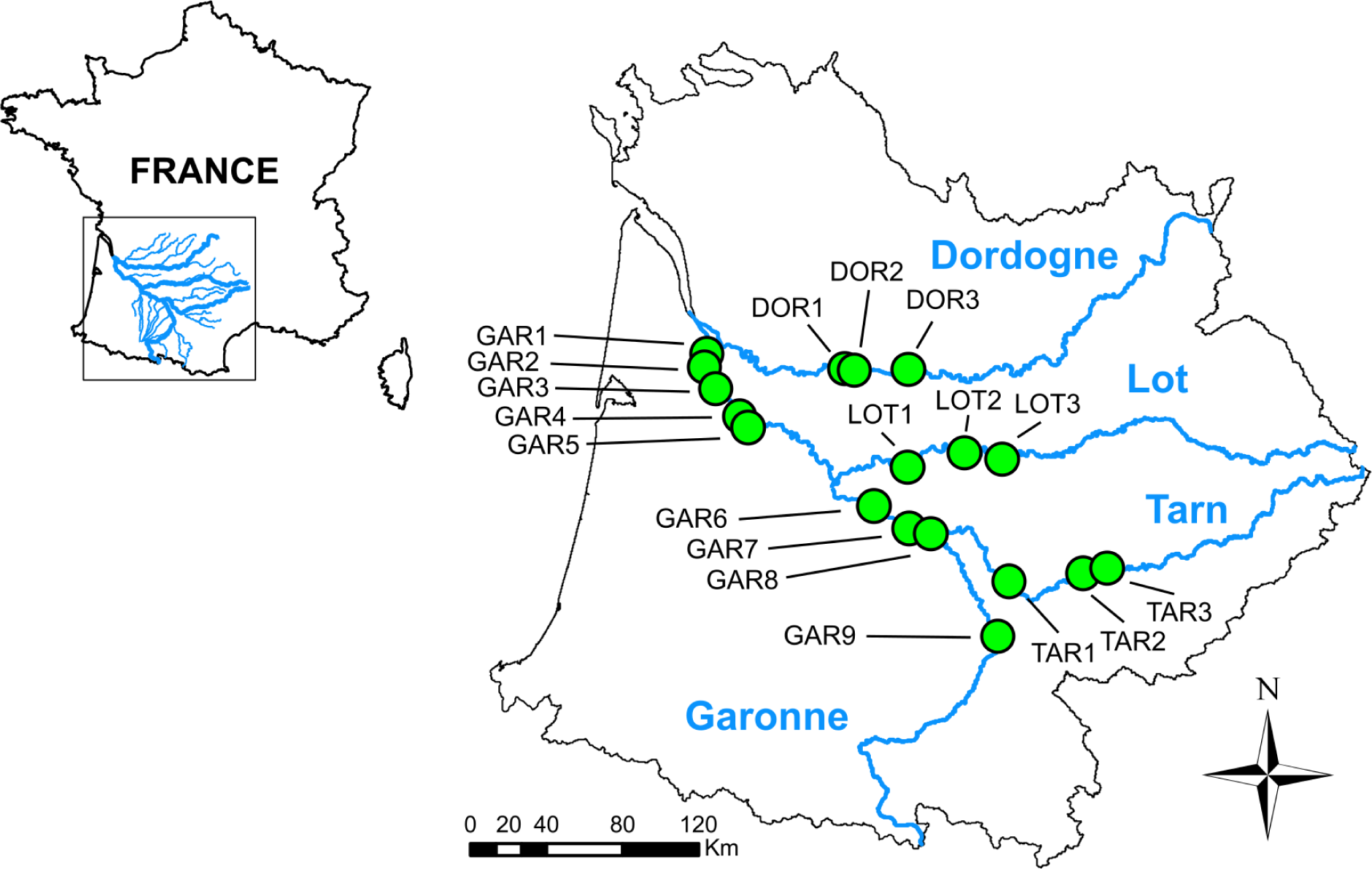
Map of Southwestern France representing the four sampled rivers (Garonne, Dordogne, Lot and Tarn) and the different sites where European catfish individuals have been sampled (green dots).

**TABLE 1:** Information on *Silurus glanis* sampling sites, methods and contributors. MIGADO is a local environmental association; FFPPMAs are departmental federations of angling associations. LAB indicates that the sampling has been carried out by laboratory members.

| Site Code | Sampling contributor | Sampling method | Year | River | Site | Longitude | Latitude | Distance to the river mouth (km) | N |
| --- | --- | --- | --- | --- | --- | --- | --- | --- | --- |
| DOR1 | Pascal Verdeyroux | Angling | 2016-2017 | Dordogne | Lamonzie / Prigonrieux | N 44°84'96.8" | E 0°38'14.4" | 148.125 | 15 |
| DOR2 | Pascal Verdeyroux | Angling | 2016-2017 | Dordogne | Bergerac | N 44°84'60.1" | E 0°45'00.3" | 155.073 | 12 |
| DOR3 | Pascal Verdeyroux | Angling | 2016-2017 | Dordogne | Mauzac | N 44°86'01.6" | E 0°80'41.1" | 185.991 | 96 |
| GAR1 | Professional fishermen | Fyke netting | 2016 | Garonne | Bassens | N 44°89'37.4" | O -0°60'92.9" | 9.000 | 30 |
| GAR2 | Professional fishermen | Fyke netting | 2016 | Garonne | Saint-Jean / Bouliac | N 44°83'09.3" | O -0°55'05.8" | 17.942 | 30 |
| GAR3 | Professional fishermen | Fyke netting | 2016-2017 | Garonne | Isle-Saint-Georges / Cambes | N 44°73'18.3" | O -0°47'14.4" | 32.097 | 50 |
| GAR4 | Professional fishermen | Fyke netting | 2016-2017 | Garonne | Barsac | N 44°60'82.6" | O -0°30'41.1" | 55.287 | 54 |
| GAR5 | Professional fishermen | Fyke netting | 2016-2017 | Garonne | Langon | N 44°55'78.2" | O -0°24'44.9" | 63.004 | 56 |
| GAR6 | Recreational fishermen | Angling | 2017 | Garonne | Agen | N 44°20'78.3" | E 0°60'55.7" | 174.442 | 9 |
| GAR7 | MIGADO | Trapping | 2016-2017 | Garonne | Golfech | N 44°10'54.9" | E 0°83'90.8" | 201.644 | 63 |
| GAR8 | LAB | Electrofishing | 2016 | Garonne | Malause | N 44°08'26.8" | E 0°98'47.3" | 215.556 | 17 |
| GAR9 | LAB | Electrofishing | 2016 | Garonne | Toulouse | N 43°60'06.7" | E 1°43'69.6" | 301.476 | 9 |
| LOT1 | FFPPMA 47 | Angling | 2018 | Lot | Penne d'Agenais | N 44°39'51.1" | E 0°81'77.7" | 200.528 | 11 |
| LOT2 | FFPPMA 47 | Angling | 2018 | Lot | Prayssac | N 44°46'74.6" | E 1°19'12.9" | 251.963 | 9 |
| LOT3 | LAB - FFPPMA 46 | Electrofishing / Angling | 2016-2018 | Lot | Cahors | N 44°48'52.5" | E 1°25'38.4" | 300.849 | 30 |
| TAR1 | LAB – Recreational fishermen | Electrofishing / Angling | 2015 | Tarn | Villemur | N 43°86'38.3" | E 1°50'37.9" | 282.868 | 19 |
| TAR2 | FFPPMA 81 | Angling | 2017 | Tarn | Aiguelèze | N 43°90'89.4" | E 1°98'85.7" | 342.140 | 18 |
| TAR3 | LAB | Electrofishing | 2015 | Tarn | Albi | N 43°93'61.6" | E 2°13'14.3" | 360.486 | 31 |

*Genotyping.* We extracted genomic DNA using a modified salt-extraction protocol (Aljanabi and Martinez, 1997). Ten microsatellite loci (Krieg et al., 1999) were co-amplified through two multiplexed Polymerase Chain Reactions (PCRs). We used 5-20 ng of genomic DNA and QIAGEN® Multiplex PCR Master Mix (Qiagen, Valencia, CA, USA) to perform PCR amplification. Details on loci, primer concentrations and PCR conditions are provided in Table S1 and Chiarello et al., (2019). The genotyping was conducted on an ABI PRISM™ 3730 Automated Capillary Sequencer (Applied Biosystems, Foster City, CA, USA). The scoring of allele sizes was done using GENEMAPPER® v.4.0 (Applied Biosystems, Waltham, USA).

*Genotyping quality controls*. We determined the occurrence of null alleles and potential scoring errors with the program MICROCHECKER 2.3 (Van Oosterhout et al., 2004). We assessed departures from Hardy-Weinberg (HW) equilibrium with the ‘adegenet’ R package V2.1.1 (Jombart, 2008). The program FSTAT v2.9.3.2 (Goudet, 1995) was used to assess linkage disequilibrium among loci within sites. Levels of significance for these multiple tests were adjusted using False Discovery Rate (FDR) procedures (Benjamini and Hochberg, 1995).

### Descriptive genetic variation analyses

*Genetic diversity and structure.* We first calculated mean numbers of alleles across loci (Na) and expected heterozygosities (H_exp_) for all sampling sites with the program GENETIX v.4.05 (Belkhir et al., 1996). Given the unequal number of individuals across sites, we further calculated local allelic richness (AR; Petit et al., 1998) and private allelic richness (PA; Kalinowski, 2004) by applying the rarefaction procedures implemented in ADZE v1.0 (Szpiech et al., 2008), using a threshold of N=9 individuals for rarefaction, corresponding to the lowest per-site sampling effort. We also used FSTAT v2.9.3.2 (Goudet, 1995) to calculate pairwise Fst values between sites (Weir and Hill, 2002), from which we derived within-site genetic uniqueness values (FstUNI; Coleman et al., 2013). This metric informs to which extent a population is genetically different (or “unique”) compared with other populations. To obtain within-site Fst_UNI_ values, we calculated the average of all pairwise Fst values observed between a given site and all other sites (Paz-Vinas et al., 2018). The higher a Fst_UNI_ value is, the more a site is considered as genetically unique compared with all others.

*Spatial patterns of genetic diversity and structure.* We mapped AR, PA and Fst_UNI_ values estimated for each sampling site to visually inspect spatial patterns of genetic variation within the four sampled rivers. We then tested whether these observed patterns conform to general spatial patterns of genetic variation theoretically expected at equilibrium for riverine fish species inhabiting dendritic river ecosystems. Specifically, we tested conformity with (i) an Isolation-By-Distance pattern (IBD) (Wright, 1943; Rousset, 1997; Sexton et al., 2014), (ii) a Downstream Increase in Genetic Diversity pattern (DIGD; Paz-Vinas et al., 2015; Blanchet et al., 2020) and (iii) a “mighty headwaters” pattern (i.e. an upstream increase in genetic uniqueness; Finn et al., 2011). We hypothesized that *S. glanis* populations would have reached an equilibrium state typical of well-established riverine fish populations in the studied hydrographical basin if they displayed significant conformity to the three spatial patterns described above (i.e., IBD, DIGD and “mighty headwaters” patterns).

Conformity with an IBD pattern was tested (i) by performing a Mantel test with 1,000 permutations considering topological distances among pairs of sampling sites as the explanatory variable, and genetic differentiation among pairs of sampling sites as the dependent variable, and (ii) by building a generalized linear model (glm) considering the two variables mentioned in (i). Topological distances among sites (i.e. distance following the watercourse) were calculated using the package ‘riverdist’ v.0.15 (Tyers, 2017) of the R statistical software v. 2.5.2 (R Development Core Team, 2017). Genetic differentiation among pairs of sampling sites was measured as F_st_/(1 – F_st_) following Rousset’s recommendations (1997) for species living in linear environments like rivers, and the Mantel test was performed using the R package ‘vegan’ (Oksanen et al., 2013). We also extracted the slope value of the glm regression curve of the IBD pattern. This parameter conveys information on the dispersal capacities of the species (Comte and Olden, 2018), and was subsequently used to account for variance in dispersal distances in meta-N_e_ calculations (see Equations 1 and 2 below).

Second, we tested whether within-site AR values increase along the upstream-downstream gradient (i.e. conformity with a DIGD pattern; Paz-Vinas et al., 2015). We built a glm considering within-site AR values as the dependent variable, and the topological distance to the river mouth as the explanatory variable. We considered a Poisson error terms distribution, included a quadratic term, and calculated an R² value using the R package ‘rsq’ (Zhang, 2017).

Finally, we checked whether within-site genetic uniqueness (Fst_UNI_) increases along a downstream-to-upstream gradient (the “mighty headwaters” pattern; Finn et al., 2011). This hypothesis predicts such an upward increase in genetic uniqueness (or genetic differentiation) due to a higher isolation of upstream sites. We built a glm considering a quadratic term, a Poisson error terms distribution, topological distances of each sampling site to the river mouth as the explanatory variable and Fst_UNI_ as the dependent variable.

*Genetic clustering analyses.* We used the sparse non-negative matrix factorization algorithm (snmf; Frichot et al., 2014) implemented in the R package ‘LEA’ (Frichot and François, 2015) to (i) estimate ancestry coefficients for all sampled *S. glanis* individuals and (ii) to identify genetically-homogenous groups of individuals (i.e. clusters) at the hydrographical basin scale. This method determines the most likely number K of clusters contained in a genetic dataset without using *a priori* information concerning individuals’ origin. Once the correct number of clusters K is determined, the method assigns each individual in the dataset to one cluster based on its proportions of genetic ancestry to such cluster (i.e. individuals’ ancestry coefficients). We tested K values ranging from K=1 to K=20. We ran 1,000 replicates for each tested K value and determined which K value and which replicate (among the 20,000 replicates we ran) best explained our genotypic data using the cross-entropy criterion implemented in ‘LEA’.

### Estimation of *Silurus glanis* effective population sizes, abundances, and densities

We aimed to estimate the effective population sizes (N_e_), abundances (N_c_) and densities of *S. glanis* in the four main rivers of the Garonne hydrographic basin by accounting for the connectivity among these four rivers, the spatial structure of current meta-populations, and the complex demographic history observed for this species in the study area (e.g. by considering recent introduction events from single or multiple sources, and an ongoing demographic expansion process). We undertook a five-step process to meet this target (summarized in Figure 2).

**FIGURE 2.**
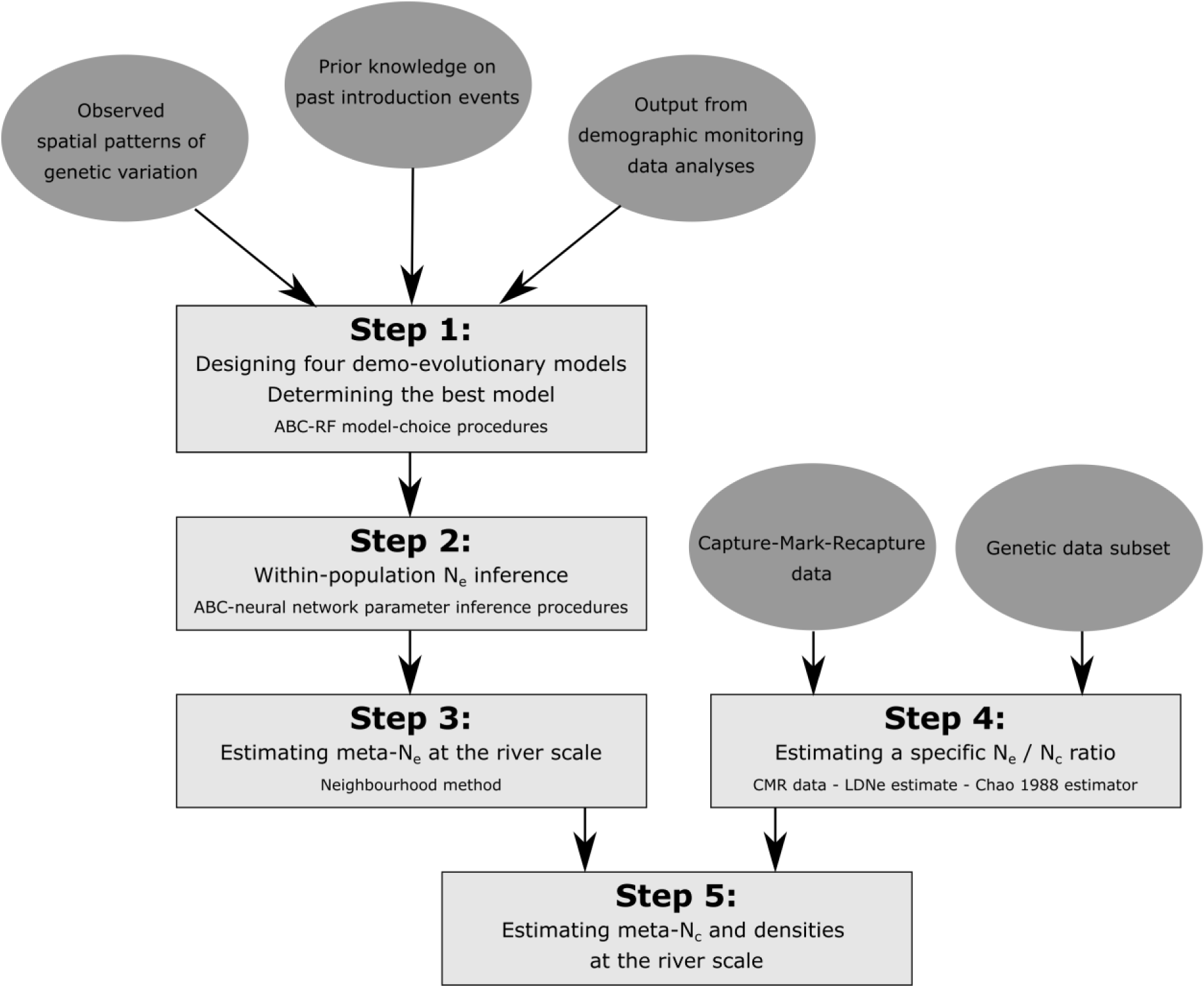
Workflow describing the five different steps we made to determine *Silurus glanis*’ metapopulation densities and effective (meta-Ne) and census (meta-Nc) sizes.

*Step 1: Approximate Bayesian Computations – Random forest (ABC-RF) procedures for model-choice (Figure 2).* In population genetics, ABC algorithms are generally used to approximate posterior distributions of model parameters for complex demographic and evolutionary (hereafter, “demo-evolutionary”) models for which the likelihood function is intractable. This is done by comparing genetic data simulated under these models with empirical genetic data (Beaumont, 2010; Csilléry et al., 2010). This comparison is done indirectly, through the use of summary statistics of genetic variation.

We designed four demo-evolutionary models (Figure 3). The first model (ONE_SYM, for “One source & Symmetric gene flow”; Figure 3A) considers that 18 demes (corresponding to the 18 groups of individuals sampled at each sampling site, see Table 1) have been founded by individuals originating from a single ancestral population of unknown origin (Source 1 in Figure 3A) 11 generations before our sampling took place (i.e. in 1983, date of the first known successful introductions in the Garonne-Dordogne river basin, considering a generation time of 3 years; Kottelat and Freyhof, 2007). Following the introduction, all demes underwent a demographic expansion process until they reached their current effective sizes (Figure 3), which could vary from 5 to 5,000 (bounds of the uniform prior distribution for effective size parameters). The demographic expansion process was parametrized using output from demographic data analyses (Appendix A1). Specifically, we analyzed data from a yearly standardized demographic monitoring survey conducted by the French Office for Biodiversity (Irz et al., 2022; Poulet et al., 2011), and used the slope of a regression curve between the number of captured individuals and sampling year to define a population growth parameter that was subsequently used in our simulations (see Appendix A1 for details). All demes exchanged individuals with neighbouring demes according to a linear stepping stone model (Kimura and Weiss, 1964; Paz-Vinas et al., 2013; Paz-Vinas et al., 2015), and assuming symmetric migration between pairs of demes (i.e. identical downstream- and upstream-directed migration rates, see Figure 3A). Migration rates among demes were drawn from an uniform prior distribution ranging from 0.001 to 0.01.

**FIGURE 3.**
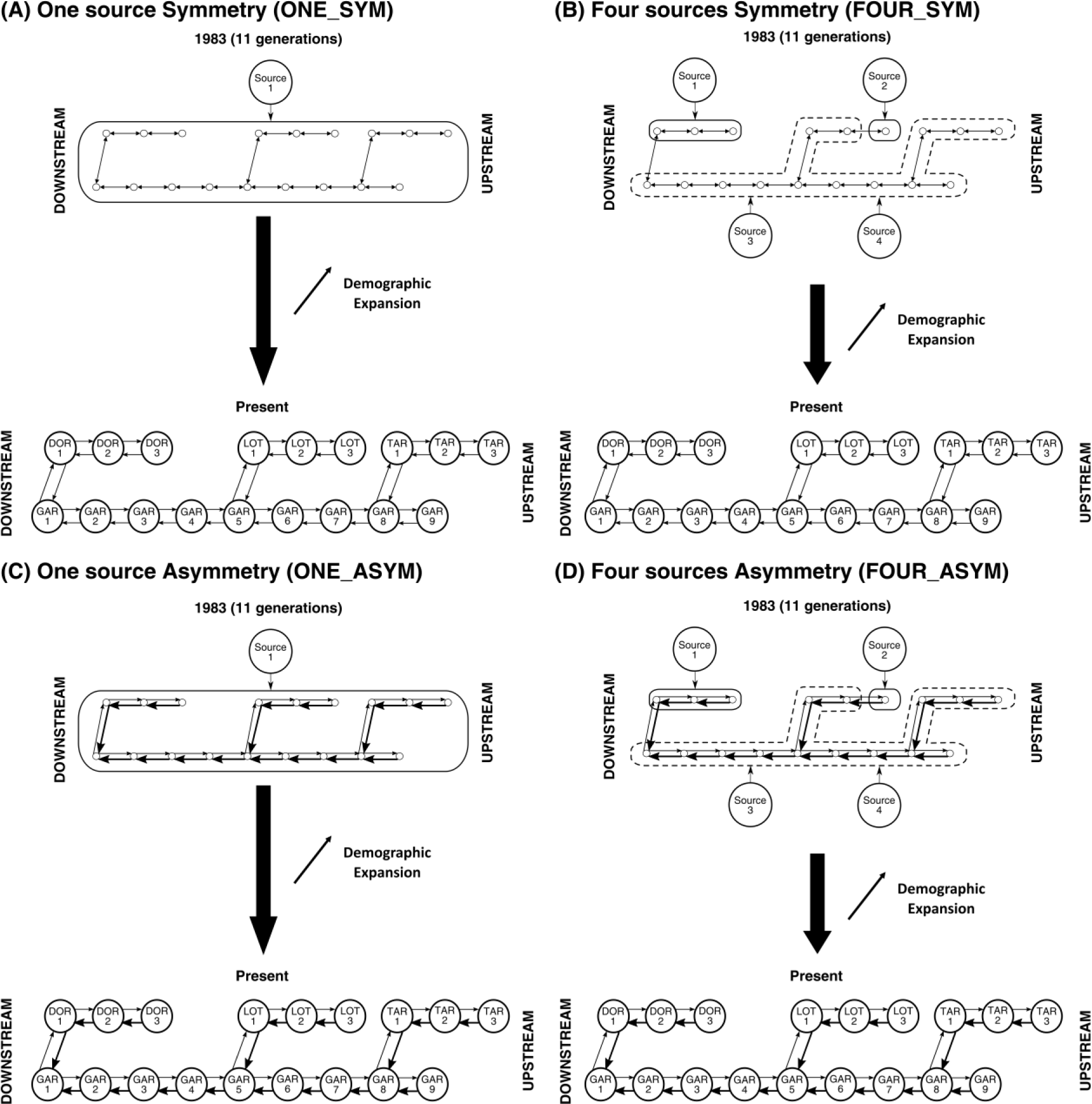
Figure representing the four competing demo-evolutionary models considered for simulating genetic data for subsequent approximate Bayesian computation analyses. Rounds represent sampling sites or putative source populations. Arrows represent inter-population gene flow

The second scenario (FOUR_SYM for “Four sources & Symmetric gene flow”; Figure 3B) stipulated that the 18 demes were the result of introductions of individuals originating from four different source populations from unknown locations (Source 1 to 4 in Figure 3B), corresponding to the K=4 number of clusters previously determined by the ‘LEA’ analysis (see Results section). We assumed that each of the four clusters was representative of a distinct genetic origin. Specifically, we considered that (i) Source 1 (Figure 3B) was the ancestral population of all individuals introduced in the Dordogne River 11 generations ago, (ii) Source 2 was the ancestral population of all individuals introduced in the most upstream site sampled in the Lot River (LOT3; Figure 3B), and (iii) individuals from Garonne, Tarn and downstream sites of the Lot (LOT1 and LOT2) originated from two different source populations (Sources 3 and 4; Figure 3B). The effective size of each source was set to be drawn from uniform prior distributions bounded between 1,000 and 10,000 individuals.

We also considered two alternative scenarios: ONE_ASYM and FOUR_ASYM (Figure 3C-D). These two scenarios were identical to ONE_SYM and FOUR_SYM except that we considered asymmetrical (i.e. downstream-biased) migration rates, a particular migration pattern expected for many riverine species due to water flow unidirectionality (Blanchet et al., 2020; Fraser et al., 2004; Kawecki and Holt, 2002; Morrissey and de Kerckhove, 2009; I. Paz-Vinas et al., 2013; Paz-Vinas et al., 2015). Downstram-directed migration rates were drawn from uniform distributions with values ranging between 0.001 to 0.01, while upstream-directed migration rates were equal to downstream-directed migration rates divided by an asymmetry parameter varying from 1.1 to 20.

We simulated genetic datasets under the four demo-evolutionary models described above by (i) sampling parameter values for a given scenario from prior distributions, (ii) simulating a genetic dataset by considering the parameter values selected in (i), (iii) calculating summary statistics from the genetic dataset simulated in (ii), and (iv) repeating steps (i) to (iii) 150,000 times *per* scenario to create a large reference table containing both the selected parameter values and the calculated summary statistics. We implemented this algorithm in a computational pipeline adapted from Rey et al. (2015) that combines diverse population genetics and statistical programs, including abcsampler (Wegmann et al., 2010), Simcoal v2.1.2 (Laval and Excoffier, 2004), PGDSpider v2.1.1.5 (Lischer and Excoffier, 2011), arlsumstat (Excoffier and Lischer, 2010), ADZE, and R (see Appendix A2 for a detailed description of the computational pipeline). For each scenario, we simulated (using Simcoal 2) 150,000 genotypic datasets based on 10 independent microsatellite markers evolving under a Generalized Stepwise Mutation Model (GSM; Kimmel and Chakraborty, 1996). Each microsatellite marker was characterized by a mean neutral mutation rate MSAT_MUTRATE_ derived from a uniform distribution. For each deme, we sampled a number of diploid individuals identical to the number of successfully genotyped individuals in our empirical dataset to ensure that the summary statistics of genetic variation calculated from our simulations were comparable with those calculated from our empirical dataset. We then computed a series of summary statistics of genetic variation for each dataset to create a reference table. In total, we calculated up to 458 summary statistics for each simulated dataset (including all pairwise Fst comparisons, means and standard deviations at the deme level for many statistics like heterozygosity, AR and PA, and global statistics calculated at the putative river basin level).

We then used the ABC-RF method (Pudlo et al., 2016) to determine which of the four demo-evolutionary models we tested best explains our empirical data (Figure 2). We applied this procedure using the R package ‘abcrf’ (Pudlo et al., 2016), by growing a forest composed of 2,000 classification trees using the 150,000 simulations *per* model we previously generated. We then assessed the robustness of the model selection procedure using OOB simulations (i.e. Out-Of-Bag simulations that were not used to grow a tree in the forest) to generate an OOB confusion matrix (Table S2) from which we extracted an OOB-based prior classification error rate. Finally, we used the Random Forest and the set of summary statistics calculated from our empirical dataset to predict which model best explains our observed data and the patterns of genetic variation observed for *S. glanis* in the Garonne-Dordogne hydrographic basin.

*Step 2: Within-population inference of effective population sizes (Figure 2).* We used the ABC parameter inference method based on neural networks implemented in the R package ‘abc’ (Blum et al., 2013; Csilléry et al., 2012) to estimate the N_e_ of the 18 demes from data simulated under the best-supported model (i.e. the FOUR_SYM model, see Results section). To obtain accurate N_e_ estimates, we generated 850,000 additional simulations under this model, which were added to the 150,000 already used for the ABC-RF model-choice procedure. We considered a subset of the most informative summary statistics (i.e. statistics displaying the highest variable importance) in the ABC-RF model-choice procedure to perform this analysis.

*Step 3. Estimating meta-N_e_ values at the river scale (Figure 2)*. We applied the Neighborhood method (Wright, 1946; Maruyama, 1971; Gomez-Uchida et al., 2013) to calculate meta-N_e_ values for each river in the Garonne-Dordogne river basin using within-deme N_e_ estimates obtained through the ABC parameter inference procedure. We considered that each river represented a meta-population, except for the Lot river, for which we considered two meta-populations (owing to the results of the genetic clustering approach and the presence of a dam): one formed by all populations located downstream of the Cahors hydroelectric dam (hereafter *Lot downstream*; including LOT1 and LOT2; Figure 1), and the other formed by the population located upstream the dam (*Lot upstream*; LOT3 in Figure 1). The Neighborhood method is suitable for linear-like environments such as rivers, as it takes into account the length of linear habitat occupied by the species and its dispersal capacities. Meta-N_e_ values were specifically calculated using the following formula (Equation 1):

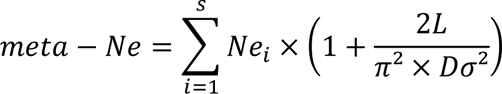

with:

*S* = Number of demes composing the meta-population

*i* = Deme *i*

*Ne_i_* = Effective size of the deme *i*

*L* = Length of the linear habitat occupied by the meta-population

*D* = Sum of N_e_ values from all demes divided by *L*

*σ*² = Dispersal parameter estimated from the slope of the IBD pattern regression curve (see Figure S2C) using the following equation (Equation 2):

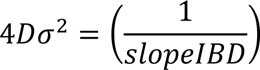

*Step 4: Estimating a specific N_e_/N_c_ ratio for S. glanis (Figure 2).* Given the lack of studies reporting N_e_/N_c_ ratios for *S. glanis*, we calculated a specific ratio (along with its 95% confidence interval) for this species using repurposed CMR data from a survey performed between 2012 and 2016 in the Dordogne River to estimate N_c_, and (ii) a subset of genotyped individuals sampled in the same river section within the same period for estimating N_e_. A detailed description of these calculations is provided in Appendix A3.

*Step 5: Estimating meta-N_c_ values and S. glanis densities at the river scale (Figure 2).* Finally, we used the N_e_/N_c_ ratio calculated in Step 4 to estimate meta-population census sizes (meta-N_c_) for the four *S. glanis* meta-populations considered (Garonne, Tarn, Lot downstream, and Dordogne), by multiplying meta-N_e_ values obtained in *Step 3* by the specific N_e_/N_c_ relationship we estimated in *Step 4* (Figure 2) For Lot upstream, a simple N_c_ estimate was calculated by multiplying the N_e_ value inferred for the LOT3 deme. Alternatively, as a comparison basis, we also calculated meta-N_c_ values using the consensual relationship N_e_/N_c_ = 0.1. We subsequently estimated average *S. glanis* densities in meta-populations by dividing obtained meta-N_c_ values (expressed as numbers of individuals) by the total area of habitat (in hectares) occupied by these meta-populations. These areas were calculated by multiplying the topological length of the river sections occupied by the meta-populations by the mean width of these river sections. Mean widths were calculated using ArcGIS v10.2 (ESRI, Redlands, CA, USA) and data from the French Theoretical Hydrographical network “RHT” (Pella et al., 2012). Topological lengths of river sections were calculated using the R package ‘riverdist’ (Tyers 2017).

## RESULTS

*Genotyping quality controls*. We found significant homozygote excess (suggestive of the presence of null alleles) for 9 locus/site pairs out of 180 possible pairs: three pairs involved the Sgl310 locus (for GAR4, GAR7 and DOR3 sites), two pairs involved Sgl325 (GAR4 and GAR7), two other pairs for Sgl5F (GAR1 and GAR3), one pair for Sgl7e (GAR5) and one pair for Sgl7159 (GAR9). We also found significant deviations from HW for two locus/site pairs (involving the locus Sgl310 for DOR3 and LOT3 sites). Given that these deviations were not generalized at both marker and sampling site levels, we retained all ten microsatellite markers for subsequent genetic analyses. We did not detect a significant linkage disequilibrium between pairs of loci across any sites.

*Spatial patterns of genetic variation.* Sampling sites displaying the highest mean allelic richness were DOR1, DOR2 and LOT1 (AR of 4.208, 4.180 and 4.020 respectively, Figure 4A; Table S3), while LOT2 and LOT3 showed the lowest mean AR (3.276 and 3.091; Figure 3A; Table S3). Mean private allelic richness was highest in LOT1 and LOT2 (PA of 0.133 and 0.106, respectively; Figure 4B; Table S3), followed by GAR9 and GAR6 (0.096 and 0.086 respectively). The lowest PA values were observed for TAR3 (0.007) and GAR8 (0.016; Figure 4B and Table S3). Similar to AR, DOR1, DOR2 and LOT1 displayed the highest Hexp values (0.715, 0.698 and 0.689 respectively, Figure 4C; Table S3), while the lowest values were found for GAR6 and LOT3 (0.596 and 0.604 respectively, Figure 4C; Table S3). The highest Fst_UNI_ values were observed in LOT3 (0.163) and LOT2 (0.128; Figure 4D and Table S3), while the lowest values were observed in GAR4 and GAR8 (0.035; Figure 4D and Table S3).

**FIGURE 4.**
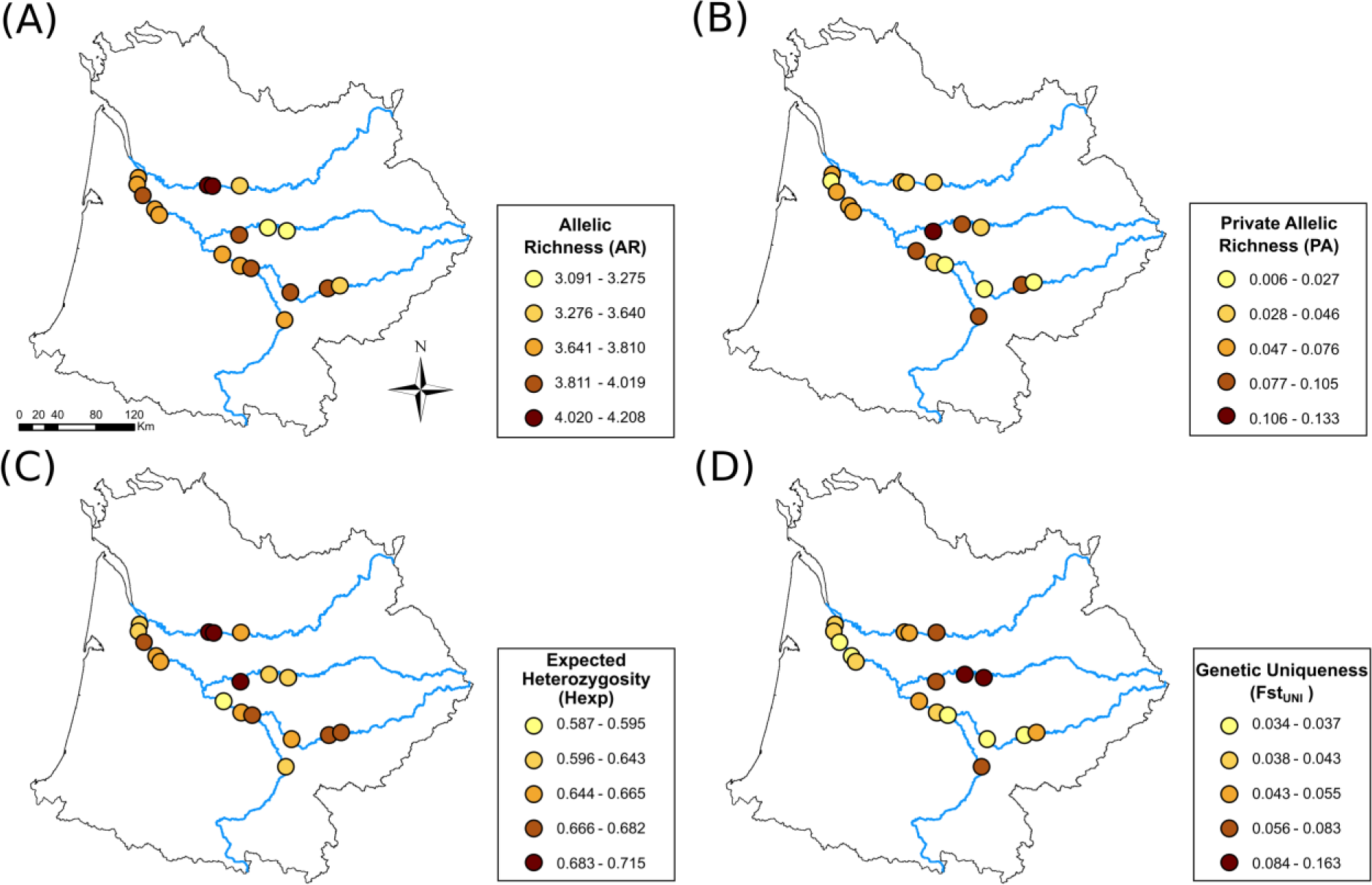
Spatial distribution of Allelic Richness (AR; A), private allelic richness (PA; B), Expected heterozygosity (Hexp; C) and genetic uniqueness (Fst_UNI_; D) for *Silurus glanis*.

We did not detect either a significant DIGD pattern or a significant “mighty headwaters” pattern (Figure S1A-B). However, we found a significant IBD pattern in *S. glanis* populations in the Garonne basin (Mantel test *r* = 0.338; p-value = 0.00089; Figure S1C). The value of the slope of the glm regression for the IBD pattern (i.e. a requisite for Equation 2 above) was equal to 1.868 x 10^-4^.

Only one of the three expected theoretical spatial patterns was verified (i.e., IBD pattern). Overall, these results suggest that European catfish meta-populations might have not yet reached genetic equilibrium at the river basin scale.

*Genetic clustering analyses.* We detected the occurrence of K=4 clusters in our genotypic dataset (Figure S2). The first cluster (cluster 1 in green in Figure 5) grouped individuals almost exclusively belonging to sites from the Dordogne River (individuals with a proportion of ancestry to cluster 1 greater than 0.8; Figure 5). A second cluster (cluster 2 in blue in Figure 5) almost exclusively grouped individuals sampled in the most upstream site from the Lot River (LOT3; Figure 5), suggesting isolation of these individuals in relation to those present in LOT1 and LOT2. The two downstream sites from the Lot River and all sites from the Tarn and Garonne Rivers were formed by a mixture of individuals belonging to the remaining two clusters (cluster 3 in yellow and cluster 4 in red; Figure 5). It is noteworthy that a majority of individuals belonging to cluster 4 were located in the two most downstream populations of the Lot River.

**FIGURE 5.**
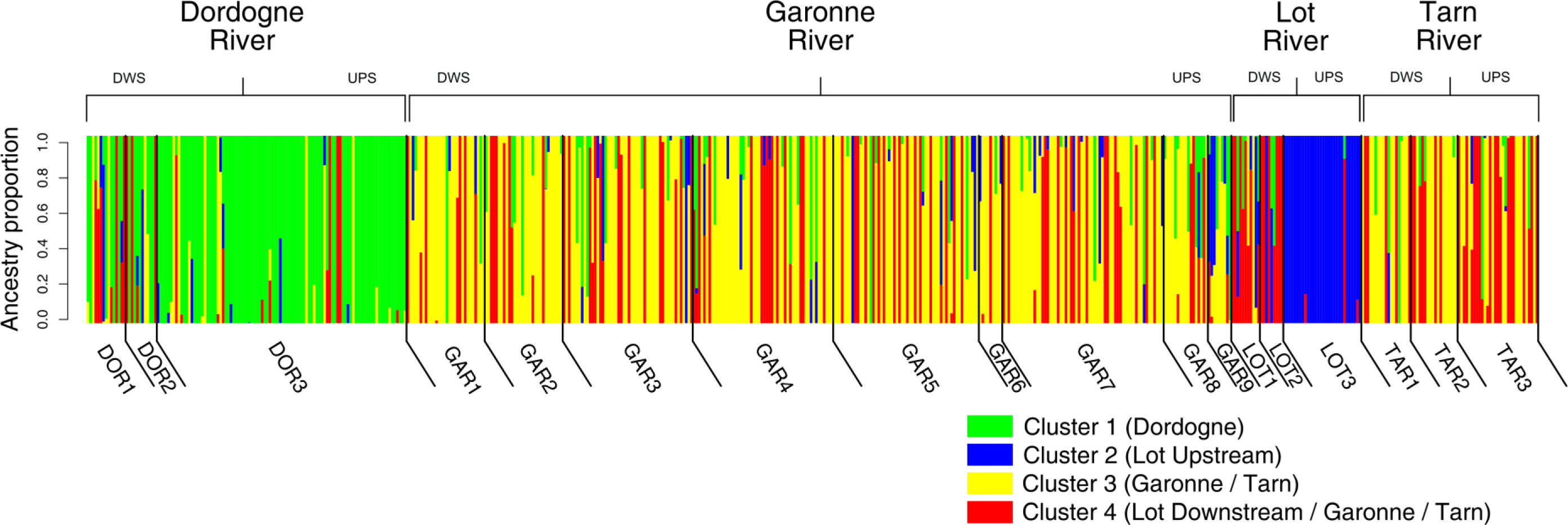
Ancestry proportions of individuals to each cluster inferred by the ‘LEA’ analysis. Individuals are sorted by River, then by following a downstream-upstream gradient. DWS = Downstream; UPS = Upstream.

### Estimation of *S. glanis* effective population sizes, abundances, and densities

*Step 1.* The ABC-RF model choice procedure indicated that the FOUR_SOURCES_SYM model was the most plausible of the four demo-evolutionary models we considered, with an estimated posterior probability of 46.8 %. Overall, we obtained a prior OOB classification error rate of 28.5% (Table S2). The highest proportions of misclassifications were observed between ONE_SOURCE_SYM and ONE_SOURCE_ASYM models (42.93 - 44.43% misclassification rates respectively; Table S2). The high number of misclassifications between these two models was likely due to the huge similarity between these models, which differed only by their migration patterns.

*Step 2.* Within-deme N_e_ values inferred through the ABC neural network approach ranged between 245 and 4,370 individuals (Figure 6). The lowest N_e_ values were found for the most upstream demes from the Lot and Dordogne rivers (LOT3 and DOR3, with N_e_ = 245 and 299 respectively; Figure 6), followed by the most downstream demes from the Lot River (LOT1, with N_e_ = 336; Figure 6). The highest values were found for two demes from the Garonne River (GAR5 and GAR1, with N_e_ = 4,370 and 4,179 respectively; Figure 6), followed by the two central demes of the Dordogne and Lot rivers (DOR2 and LOT2, with N_e_ = 3,978 and 3,964 respectively; Figure 6). The N_e_ of the four different source populations ranged between 1,592 (Source 3; see Figure 6) and 8,022 (Source 1; Figure 6).

**FIGURE 6.**
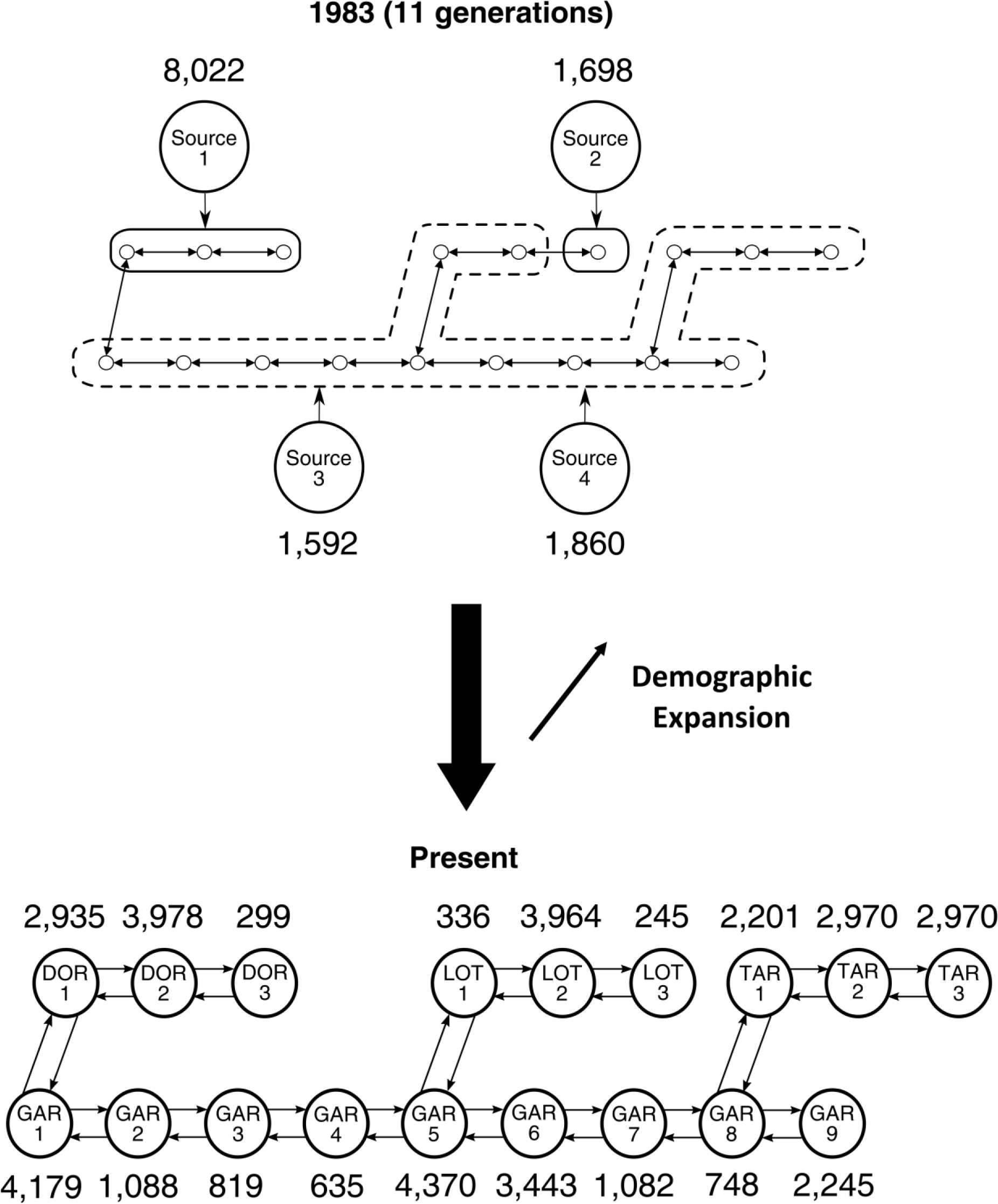
Point Ne estimates obtained under the best-supported demo-evolutionary model.

*Steps 3 to 5*. Meta-N_e_ values ranged between 245 (Lot upstream) and 19,502 (Garonne river; Table 2). Lot downstream, Dordogne and Tarn meta-populations showed intermediate meta-Ne values (4,377, 7,390 and 8,314 respectively; Table 2). The specific N_e_/N_c_ ratio we calculated using data from the Dordogne was equal to 0.3697 (CI95% = [0.2048 – 0.615]; Appendix A3). Accordingly, the meta-N_c_ of the five considered meta-populations varied between 664 [399 – 1199] and 51,847 [31,186 – 93,615] individuals for the Lot upstream and Garonne meta-populations respectively (Table 2), for a total estimated number of individuals of 106,828 [64,255 – 191,681] across all four rivers. The Tarn River meta-population was the one with the highest average *S. glanis* population density (i.e. 19.093 [11.484 – 34.466] individuals/hectare; see Table 2), while the Lot upstream population displayed the lowest densities of *S. glanis* individuals (2.664 [1.603 – 4.810] individuals/hectare; Table 2). The consensual N_e_/N_c_ ratio led to higher estimates of meta-N_c_ and density overall (Table 2).

**TABLE 2:** Meta-N_e_, Meta-N_c_ and density estimates for the five meta-populations.

| Meta-population | Linear distance (km) | Mean Width (m) | Meta-$N_e$ | Using a specific $N_e/N_c$ ratio = 0.3697 [0.2048 – 0.615] | | Using a consensual $N_e/N_c$ = 0.1 ratio | |
| --- | --- | --- | --- | --- | --- | --- | --- |
| | | | | Meta-$N_c$<br>(number of individuals) | Density<br>(individuals / hectare) | Meta-$N_c$<br>(number of individuals) | Density<br>(individuals / hectare) |
| Dordogne | 162.325 | 106.388 | 7 390.348 | 19 989.670 [12 023.779 – 36 093.238] | 11.575 [6.962 – 20.900] | 73 903.475 | 42.794 |
| Lot Downstream | 119.073 | 79.68 | 4 377.079 | 11 839.278 [7 121.321 – 21 376.935] | 12.479 [7.506 – 22.531] | 43 770.797 | 46.134 |
| Lot Upstream | 29.12 | 85.652 | 245.669 | 664.497 [399.695 – 1199.811] | 2.664 [1.603 – 4.810] | 2 456.670 | 9.849 |
| Garonne | 316.141 | 150.61 | 19 502.559 | 51 847.537 [31 186.274 – 93 615.626] | 10.889 [6.550 – 19.661] | 195 025.591 | 40.959 |
| Tarn | 139.760 | 84.28 | 8 314.023 | 22 489.434 [13 526.916 – 40 597.377] | 19.093 [11.484 – 34.466] | 83 140.235 | 70.584 |

## DISCUSSION

In this study, we characterized the spatial patterns of genetic variation of a non-native large-bodied freshwater fish (i.e. the European catfish *Silurus glanis*) in four large rivers of the Garonne-Dordogne hydrographical basin (Southwestern France). The observed patterns of genetic variation were subsequently combined with prior knowledge of past introductions (e.g. date of first known successful introductions in the basin) and with output from demographic monitoring analyses to define, parametrize and calibrate competing demo-evolutionary models. We then (i) determined using ABC-RF model-choice procedures the demo-evolutionary model – among a set of four competing models – best explaining the observed patterns of genetic variation, and (ii) used ABC parameter inference procedures on data simulated under the best-supported model and a species-specific N_e_/N_c_ ratio to (iii) determine densities, effective and census sizes observed for five *S. glanis* meta-populations in the Garonne-Dordogne hydrographical basin. Overall, our study shows how combining snapshot population genetic surveys, demographic analyses, prior knowledge of introduction events and powerful inferential frameworks could help estimate both effective and census population sizes for non-equilibrium meta-populations of hard-to-monitor species.

### Genetic variation of Silurus glanis across the Garonne-Dordogne hydrographical basin

Descriptive genetic analyses revealed moderately high genetic diversity levels for *S. glanis* across the Garonne-Dordogne hydrographical basin, with values for genetic indices (e.g. allelic richness) akin to those estimated for other common freshwater fish species native to this hydrographical basin (e.g. *Squalius cephalus*, *Leuciscus burdigalensis,* or *Barbatula barbatula;* (Fourtune et al., 2016; Paz-Vinas et al., 2018). Interestingly, the observed levels of genetic diversity were also similar to those found in other *S. glanis* native populations from Central and Eastern Europe using a comparable set of microsatellite loci (Castagné et al., 2023; Triantafyllidis et al., 2002), although they contrast with the small genetic diversity levels recently observed in Northern Europe peripheral native populations (e.g. Swedish populations; Palm et al., 2019). Overall, our population genetic assessment suggests that significant genetic variation has been introduced and retained over time in *S. glanis* populations in the Garonne hydrographical basin since the first known introduction event in 1983. The considerable levels of genetic diversity and the high effective meta-population sizes observed in *S. glanis* populations at the hydrographical basin scale could be the result of multiple introduction events involving individuals from different genetic sources, as is the case for other aquatic invasive species inhabiting the study area (e.g. the spiny-cheek crayfish *Procambarus clarkii*; Paz-Vinas et al., 2021). This was also (i) suggested by a previous study having conducted a broader genetic survey at the European scale (Castagné et al., 2023), (ii) evidenced by our genetic clustering analyses, which revealed the presence of four genetically distinct groups of individuals (clusters), and (iii) confirmed by the ABC-RF model-choice procedure we conducted, which provided higher support for demo-evolutionary models involving four genetically-distinct sources of individuals instead of one single source.

Other concomitant factors may have contributed to the maintenance of much – if not all – of the genetic variation introduced in the basin over time, including the very long timespan of *S. glanis* individuals (i.e. up to 80 years; Kottelat and Freyhof, 2007), their high behavioural plasticity (Cucherousset et al., 2018), the rapid demographic expansion this species has experienced (Poulet et al., 2011; see also appendix A1), its low interest as a source of human food in France and the popular practice of catch-and-release angling by trophy anglers targeting this species (which limits the impact of recreational angling on the demographic regulation of this species by individuals’ removal (Arlinghaus, 2007; Cucherousset et al., 2018).

Concerning the spatial patterns of genetic variation, we did not detect significant DIGD or “mighty headwaters” patterns like those found for other common native freshwater fish species in-habiting the Garonne hydrographical basin scale (Fourtune et al., 2016; Paz-Vinas et al., 2018), although we detected a significant pattern of isolation by distance. This suggests that *S. glanis* populations might not have reached migration / genetic drift equilibrium, and we hypothesize that the species cannot yet be considered fully established at the hydrographical basin scale. The species might still be experiencing an expansion phase, as suggested by the analysis of data from the yearly national demographic surveys used to parametrize the simulations (Appendix A1). It is noteworthy that even demographically stable populations might be observed long before populations reach genetic equilibria states (e.g. migration/drift equilibrium) or display theoretically expected spatial patterns of genetic variation, as there are generally lag times between ecological and demographic events and their associated genetic consequences (Landguth et al., 2010; Epps and Keyghobadi, 2015; Gargiulo et al., 2024). We suppose that these lag times could be long (in calendar years) for species with long generation times and/or timespan like *S. glanis*.

### Selecting the most plausible demo-evolutionary model

Our snapshot genetic assessment of *S. glanis* meta-populations was particularly useful in building demo-evolutionary models for subsequent N_e_ estimations. For instance, our genetic clustering analysis suggested the presence of four genetically distinct groups of individuals (clusters) in the study area. Two of these clusters were well-defined in space: one was composed of individuals mainly found in the Dordogne River, and the second was composed of individuals from the most upstream population in the Lot River. Individuals from the two other clusters mainly occurred in downstream populations of the Lot River and the Garonne and Tarn Rivers. Local differentiation through differential genetic drift effects across rivers or differential founder effects following independent introduction events involving individuals from the same genetic pool could be an explanation for the occurrence of these four clusters. However, the relatively low number of generations having spanned between the first known introduction event and the beginning of our sampling (i.e. 11 generations, considering a generation time of 3 years; Kottelat and Freyhof, 2007), together with the long lifespan of the species (which makes it plausible that the biggest individuals we sampled originated from the first introduction events in the 80s) and the putative information on the origin of introduced populations we gathered suggest that these four clusters result from at least four large independent introductions of individuals coming from four different and unknown genetic sources. Indeed, the individuals sampled in the Garonne, Tarn and downstream sites of the Lot rivers may originate from at least two large introduction events involving two genetically distinct source populations (corresponding to clusters 3 and 4 in Figure 5). At least one introduction event involving individuals from a third genetic source (corresponding to cluster 2 in Figure 5) may have occurred in the most upstream site in the Lot River, while introductions in the Dordogne River might have involved a fourth source genetically distinct from the three others (Cluster 1 in Figure 5).

Our ABC-RF model-choice procedure indicated that the demo-evolutionary model – among the set of four competing models we tested – that best fits with the observed genetic data involves introductions from four genetically distinct sources and symmetric migration. The model-choice procedure was robust: the prior OOB classification error rate we obtained was moderately low, similar to error rates obtained in other studies applying ABC-RF procedures in invasion genetics contexts (e.g. Fraimout et al., 2017; Javal et al., 2019), and mostly driven by high misclassifications between models differing only by their dispersal modalities (symmetric vs. asymmetric dispersal). The symmetric dispersal modality assumed by our best model contradicts general expectations (predominance of downstream-biased asymmetrical gene flow in riverine systems; Paz-Vinas et al. 2013, 2015) and agrees with observations suggesting that asymmetrical gene flow may be less frequent than generally expected in riverine species (Alther et al., 2021; Blanchet et al., 2020). Although we tried to consider as many characteristics as possible in our demo-genetic models (inter-river connexions, dispersal patterns among sites, demographic expansion, multiple introduction events), our models were not suited to assess when these introductions happened (as we considered the date of the first known introduction event as the date for all introductions) or where they happened exactly within each meta-population (as individuals were supposed to be introduced in all locations within a meta-population for each introduction event).

### Inference of Ne, of Nc, and Ne/Nc values

Here, we aimed to infer the local effective population sizes of meta-populations that severely derive from the assumptions of the Wright-Fisher model. Indeed, the monitored *S. glanis* meta-populations are structured, live in continuous linear habitats instead of isolated river bodies, have experienced demographic changes (i.e. expansion) over time, exchange migrants among them, and might have received – or have been founded by – individuals originating from different genetic sources, located outside the study area. By adopting model-based simulation procedures calibrated with prior knowledge of the invasion process, information derived from demographic analyses, and information from a snapshot genetic variability assessment, we aimed at taking into account as much as possible the complexity of our study model and ecosystem, approximating at best the biological invasion process experienced by *S. glanis* in the Garonne hydrographical basin (e.g. time since introduction events, number of potential different genetic sources, population growth parameters, migration patterns). Nonetheless, some factors that are known to affect effective population size estimation such as overlapping generations or variance in reproductive success (Gargiulo et al., 2023; Nunney, 1993; Robinson and Moyer, 2013) have not been accounted for, given that the backwards-in-time genetic data simulation tool we used does not allow simulating such specificities. The impact these deviations from the assumptions of our simulation models might have had on our N_e_ estimations is unknown. However, we are confident that the potential loss of precision due to these deviations is largely compensated by having taken into account many other model study characteristics.

To translate N_e_ and meta-N_e_ estimates to N_c_, meta-N_c_ and density estimates, we used both the consensual N_e_/N_c_ ratio of 0.1 and a population-specific ratio estimated using genetic data and repurposed Capture-Mark-Recapture (CMR) from the Dordogne river (Appendix A3). It is recommended to use taxon-specific N_e_/N_c_ ratios when available to assess N_e_ or N_c_ through a N_e_/N_c_ relationship (Laikre et al. 2021), to reduce the substantial variation that can be observed depending on the taxonomic groups and species life history traits (Laikre et al. 2021; Frankham et al. 2021). However, we provide here estimates of N_c_, meta-N_c_ and density estimates calculated with the consensual N_e_/N_c_ ratio as a comparison basis, and to set a precautionary second-level confidence interval. Indeed, the CMR study from which we repurposed the data for estimating the population-specific N_e_/N_c_ ratio was not specifically designed to estimate N_c_ (Appendix A3), and there might be some uncertainty around the *S. glanis* N_e_/N_c_ we calculated, which was equal to 0.3697 (CI95% = [0.2048 – 0.615]). Nonetheless, this value was similar to the mean value estimated for nine Bony freshwater fish (0.389) by Hoban et al. (2020), though it remains slightly higher than the consensual 0.1 value, which might probably lead to the overestimation of N_c_, meta-N_c_ and density estimates in our study (Table 2). Further work is needed to develop genetic and census-size-estimation-oriented CMR sampling designs allowing more accurate N_e_/N_c_ ratio calculations for this species, both in invasive and native populations.

## CONCLUSIONS

Our study shows how prior knowledge, demographic data, population genetic assessments, and simulation-based genetic inference procedures can be integrated into a common framework to estimate N_e_, N_c,_ and densities of individuals for non-equilibrium meta-populations having experienced recent – and complex – demographic dynamics (e.g. for invasive species’ meta-populations) and/or from species that are particularly hard to monitor using direct estimation methods. The information generated under such a framework can be of prime value for biodiversity managers, as it may simultaneously provide key information on the demographic and genetic state of populations, and on potential invasion pathways used by the studied species. We also provide a first Ne/Nc ratio estimate for *S. glanis*, information that can help assess N_e_ or N_c_ in other invasive or native populations. More specifically, the N_c_ and density estimates we obtained for *S. glanis* might help quantify the potential impact of this species on native and protected ichthyofauna and fisheries, e.g. by coupling these results with those from diet analyses (Guillerault et al., 2019, 2017), body-size spectra (Boulêtreau and Santoul, 2016) and others. This may ultimately help guide future *Silurus glanis* population control actions within the four main rivers of the Garonne-Dordogne hydrographical basin.

## Author Contributions

The study was designed by IP-V, FS and GL. IP-V wrote the manuscript with the help of GL, FS, SB and MC. FS, SB and MC collected samples. GL, CV and JF produced genetic data. IP-V conducted the population genetic and statistical analyses.

## Acknowledgements

We are very grateful to all persons and organisations who contributed to the field sampling, including laboratory members, recreational and professional anglers, MIGADO, FFPPMAs 46, 47 and 81, and more particularly Mr. Pascal Verdeyroux from EPIDOR, who also conducted the Capture-Mark-Recapture procedure. We also thank the “Génopole Toulouse” for help with genotyping, and the CALMIP group for the access to their High-Perfomance-Computing facilities (computer cluster “Eos”; allocation T18005). The Centre de Recherche sur la Biodiversité et l’Environnement was supported by “Investissement d’Avenir” grants (CEBA, ref. ANR-10-LABX-0025; TULIP, ref. ANR-10-LABX-41).

## Data accessibility and Benefit-Sharing

The genotype data using in this study, as well as the scripts to conduct the genetic data simulations for the abc-rf procedure will be deposited in dryad upon acceptance of the paper.

Benefits Generated: benefits from this research will arise from the sharing of our data upon acceptance.

