## Supplementary Tables, Figures and appendices for "Genetic-based inference of densities, effective and census sizes of expanding riverine meta-populations of an invasive large-bodied freshwater fish (*Silurus glanis L.*)"

**Table S1** : Details on the microsatellite loci used in this study, based on Krieg et al. (1999). More details on primer concentration and PCR conditions can be found in Chiarello et al. (2019)

| Locus | GenBank  Accession ID | Multiplex Kit |
| --- | --- | --- |
| Sgl310INRA | AF146414 | SilA |
| Sgl325INRA | AF146415 | SilA |
| Sgl33INRA | AF146416 | SilA |
| Sgl5fINRA | AF146418 | SilA |
| Sgl7eINRA | AF146423 | SilA |
| Sgl1154aINRA | AF146410 | SilB |
| Sgl140aINRA | AF146412 | SilB |
| Sgl695INRA | AF146419 | SilB |
| Sgl7159INRA | AF146420 | SilB |
| Sgl7fINRA | AF146425 | SilB |

**Table S2 :** OOB confusion matrix obtained from a random forest procedure applied to ABC (ABC-RF) composed of 2000 classification trees and 600,000 simulations (150,000 per model). The numbers of correct assignments (simulations generated by a model and assigned to this same model by the procedure) are represented in bold. The overall OOB classification error rate of this random forest is 28.5%.

|  | Percentage of OOB simulations assigned to models: | | | |
| --- | --- | --- | --- | --- |
| Models under which OOB simulations have been generated: | ONE_SOURCE_SYM | ONE_SOURCE_ASYM | FOUR_SOURCES_SYM | FOUR_SOURCES_ASYM |
| ONE_SOURCE_SYM | **57.04%** | 42.93% | 0% | 0.03% |
| ONE_SOURCE_ASYM | 44.43% | **55.53%** | 0% | 0.04% |
| FOUR_SOURCES_SYM | 0% | 0% | **86.2%** | 13.8% |
| FOUR_SOURCES_ASYM | 0% | 0% | 12.6% | **87.4%** |

**Table S3** : Table summarizing summary statistics of the genetic variability obtained for each sampled *Silurus glanis* population. N_ALL_ corresponds to the mean number of alleles observed across loci; He corresponds to the expected heterozygosity; AR and PA corresponds to mean allelic richness and mean private allelic richness respectively, and FstUNI corresponds to an estimate of the genetic uniqueness of each population. A minimum sample size of 11 was considered for rarefaction procedures to obtain AR and PA estimates.

| Site Code | River | N_ALL_ | He | AR | PA | Fst_UNI_ |
| --- | --- | --- | --- | --- | --- | --- |
| DOR1 | Dordogne | 5.9 | 0.715 | 4.208 | 0.068 | 0.056 |
| DOR2 | Dordogne | 5.6 | 0.698 | 4.181 | 0.037 | 0.047 |
| DOR3 | Dordogne | 7 | 0.666 | 3.641 | 0.043 | 0.084 |
| GAR1 | Garonne | 6.1 | 0.643 | 3.765 | 0.071 | 0.043 |
| GAR2 | Garonne | 5.8 | 0.637 | 3.761 | 0.024 | 0.040 |
| GAR3 | Garonne | 7.3 | 0.671 | 3.885 | 0.077 | 0.036 |
| GAR4 | Garonne | 6.5 | 0.664 | 3.811 | 0.060 | 0.035 |
| GAR5 | Garonne | 6.8 | 0.661 | 3.766 | 0.066 | 0.039 |
| GAR6 | Garonne | 5 | 0.596 | 3.755 | 0.086 | 0.048 |
| GAR7 | Garonne | 6.5 | 0.660 | 3.760 | 0.046 | 0.039 |
| GAR8 | Garonne | 5.3 | 0.669 | 3.903 | 0.016 | 0.035 |
| GAR9 | Garonne | 4.8 | 0.638 | 3.791 | 0.096 | 0.072 |
| LOT1 | Lot | 5.3 | 0.689 | 4.020 | 0.133 | 0.067 |
| LOT2 | Lot | 3.6 | 0.638 | 3.276 | 0.106 | 0.128 |
| LOT3 | Lot | 4.1 | 0.604 | 3.091 | 0.042 | 0.163 |
| TAR1 | Tarn | 5.6 | 0.652 | 3.863 | 0.028 | 0.037 |
| TAR2 | Tarn | 6 | 0.683 | 3.959 | 0.083 | 0.038 |
| TAR3 | Tarn | 5.3 | 0.672 | 3.623 | 0.007 | 0.052 |

**Appendix A1 :** Defining a parameter using temporal demographic monitoring surveys data to consider the demographic expansion observed for *Silurus glanis* in the Garonne-Dordogne river basin into Simcoal 2 simulations.

Simcoal 2 is a backwards in time genetic data simulator in which past effective deme sizes must be defined in relation to current effective deme sizes (i.e. at t = 0). We thus simulated demographic expansions (forward in time) in all the 18 Garonne-Dordogne demes by specifying historical events in which we considered that the relative effective deme sizes at t = -11 generations (the time elapsed between the first introduction of individuals in the basin and the time of sampling) were Δ times the effective deme sizes computed at t = 0 (present time), with Δ being a value smaller than 1 to simulate demographic expansion. For instance, for Δ = 0.5, a deme having an effective size of 200 individuals at t = 0 would have had a size of 100 individuals at t = -11.

To estimate Δ, we first repurposed data from a French National Demographic monitoring program based on ∼4 decades of standardised electrofishing surveys conducted in ~6000 sampling sites distributed across all hydrographic networks in mainland France rivers (Poulet et al. 2011 ; Irz et al. 2022). We extracted from the “Naïades” web portal (<https://naiades.eaufrance.fr/>) data on all *S. glanis* captures made at all sampling sites located within the Garonne-Dordogne river basin from the beginning of electrofishing surveys until 2015.

Though the first known introduction of *S. glanis* was made in 1983, the first detection of an individual of that species by the monitoring program occurred in 1995, probably because populations were not large or spatially-extended enough to be detected before by the monitoring protocol (Figure Appendix A1). Since then, the number of individuals captured at the Garonne-Dordogne river basin scale significantly increased linearly (Figure Appendix A1). Here, we assumed that the linear regression between the number of captures of *S. glanis* individuals over time made by the monitoring program is representative of the overall demographic expansion trend exhibited by the species across the whole river basin. We therefore computed the equation of a linear model fitting that data (R²=0.794 ; p<0.001), which led to Equation A1-1:

$$y=10.175x-20314.716$$

with:

*x* = the year of sampling

y = the number of captured individuals

We then assessed what could be the value of Δ using Equation A1-1, considering two values of *y* separated by a period of x_0_ – x_-11_ = 33 years (i.e. 11 generations), considering Equation A1-2:

$$\Delta=\frac{1}{y_{0}}\times y_{-11}$$

with:

Δ = the relative size of a deme at time x_-11_ compared to the size of that deme at time x_0_
y_0_ = the number of captured individuals at time x_0_

y_-11_ = the number of captured individuals at time x_-11_

We thus computed the difference in number of individuals estimated through Equation A1.2 for the year 2000 (y_-11_ = 35.284) and for 2033 (y_0_ = 371.059), leading to a Δ value of 0.09509. This value was then used in Simcoal “.par” files to simulate historical events considering a demographic expansion akin to that observed in the Garonne-Dordogne river basin.

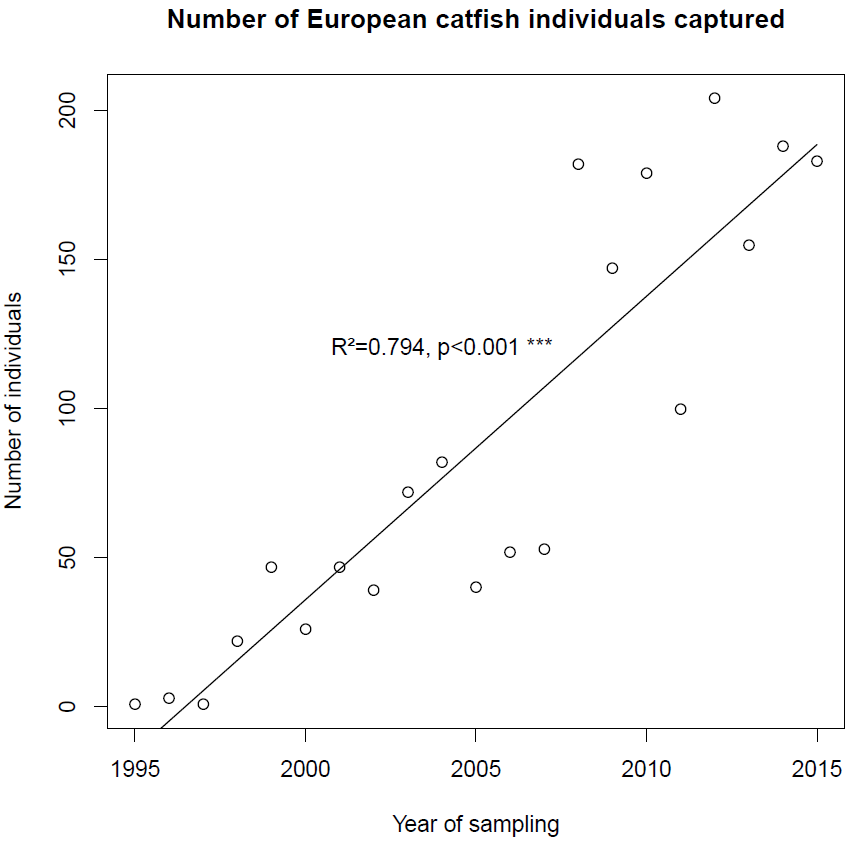

*Figure Appendix A1: Number of European catfish (Silurus glanis) individuals captured* at the Garonne-Dordogne river basin scale by a National monitoring program from 1995 to 2015. The black line represents the regression line of a linear model of equation *y* = 10.175*x* – 20314.716

**Appendix A2 :** The computational pipeline used for implementing the simulation procedure.

To implement the simulation procedure used in this study, we adapted the pipeline previously used in Rey et al. (2015) and Paz-Vinas et al. (2015) to the specificities of our study. This computational pipeline combines multiple population genetics and statistical programs (see Figure A2 for a graphical representation of the pipeline). This pipeline allowed us to (i) calculate a wide number of genetic summary statistics that cannot be obtained from only a single population genetics program, and (ii) using a flexible genetic data simulator that allowed us to account for the spatial connectivity among rivers, the demographic expansion of *Silurus glanis* observed in the Garonne-Dordogne river basin, and potential introduction events using individuals originating from population that are outside of the river system. The core of the pipeline was the program ABCsampler from the ABCtoolbox package (Wegmann et al., 2010). This program, combined to bash scripts allowed (i) automatically managing dialogs between the different programs composing the pipeline, (ii) sampling parameter values (φ_x_) from prior parameter distributions (see Materials and Methods section), and (iii) feeding the simulation program with φ_x_. We used the coalescent-based, backwards-in-time genetic data simulator SIMCOAL v2.1.2 (Laval and Excoffier, 2004) to simulate the microsatellite datasets (D_x_) for each of the four models given φ_x_. The conversion program PGDSpider v2.0 (Lischer and Excoffier, 2011) was integrated into the pipeline to convert each D_x_ generated by SIMCOAL v2.1.2 in *arlequin* format (Excoffier and Lischer, 2010) to other formats specific to the softwares used for calculating summary statistics. The software arlsumstat (Excoffier and Lischer, 2010) was fed with D_x_ in *arlequin* format to calculate over loci and for each deme expected heterozygosities (*He*), the Garza-Williamson's index (*GW*), global *F_st_,* and *F_is_* values, as well as among-demes pairwise *F_st_* values. In parallel, PGDSpider v2.0 converted the D_x_ in *arlequin* format to the *structure* format (Pritchard et al., 2000), which corresponds to the input format of the software ADZE v1.0 (Szpiech et al., 2008). This last program was used to estimate, at the deme level, the means, variance and standard deviations of allelic richness (AR) and private allelic richness (PA) among loci. Finally, we used the R statistical software (R Development core Team 2017) to extract, concatenate and store all the summary statistics in a single file.

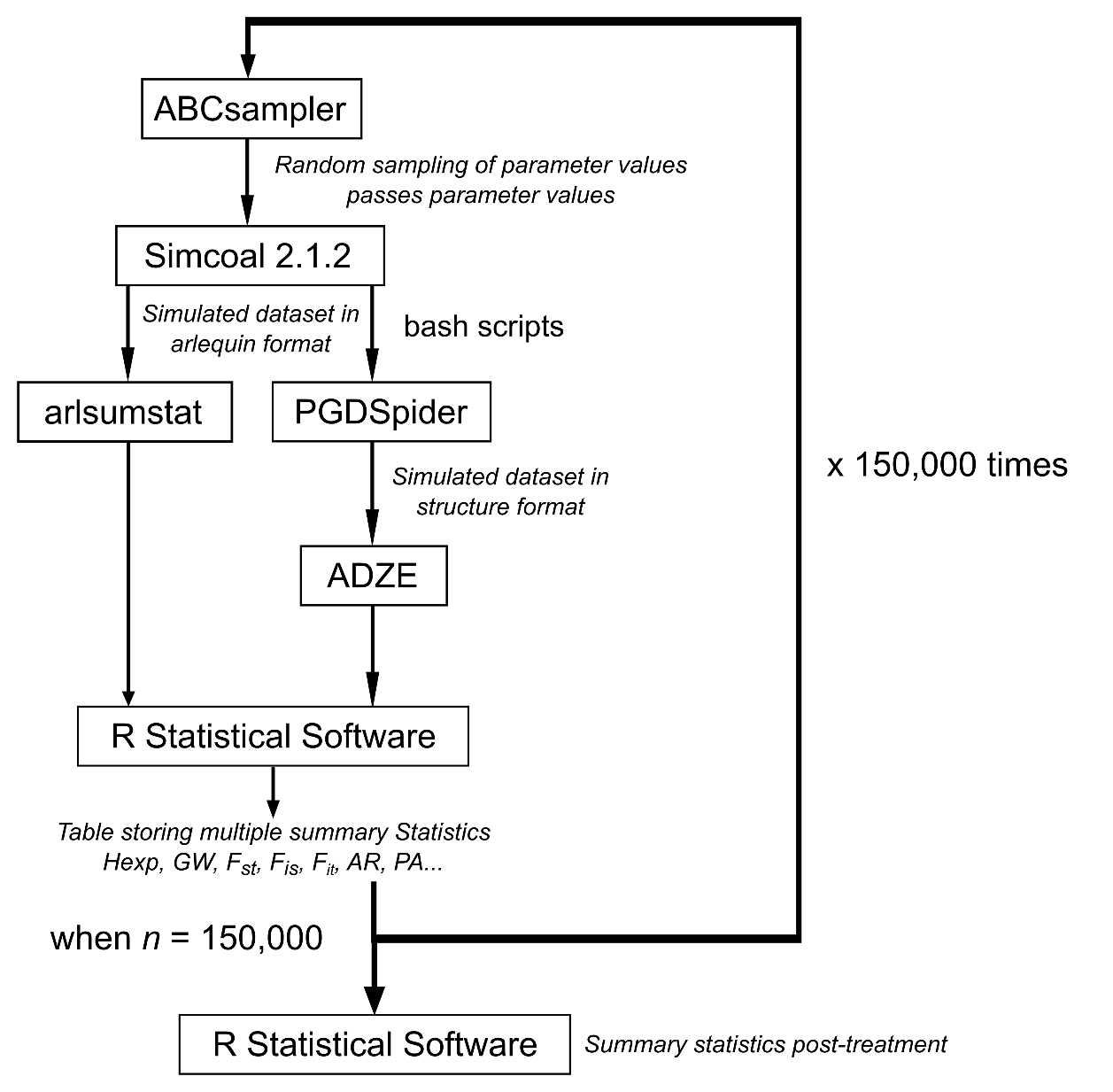

*Figure Appendix A2 :* Graphical representation of the computational pipeline of programs used for the for the serial simulation of genetic datasets for implementation of the ABC-RF algorithm.

We used CMR data compiled by EPIDOR (“Etablissement Public Territorial du Bassin de la Dordogne”) between April 2012 and July 2016 on a section of the Dordogne river that spans from Pessac to Mauzac (EPIDOR, Verdeyrou, Guerri, 2016). The CMR procedure conducted by EPIDOR was not specifically designed to calculate the N_c_ of the *S. glanis* population, but rather to characterize the spatial distribution of the species along the Dordogne River, for making biometrical measurements on individuals, and to determine their preferred habitats and diet. Therefore, some of the conditions necessary for the application of many methods aiming at estimating N_c_ from CMR data were not implemented, such as for instance having a defined temporal spacing between multiple recapture campaigns (recaptures and tagging were made continuously). Their data was nevertheless compatible with the application of the Chao point abundance estimator (Chao, 1988).

This method allows estimating the abundance of individuals in a population from CMR data even when the probability of recapture of individuals is low (Chao, 1988). The estimate of the population size N_c_ can therefore be obtained by applying the following formula:

$$Nc=S+\frac{F_{1}^{2}}{{2F}_{2}}$$

with:

S = Number of individuals captured and marked

F1 = Number of individuals captured and marked, but not subsequently recaptured

F2 = Number of individuals captured, marked and recaptured

This method, unlike other estimation methods, is conservative, and tends to underestimate the values of N_c_. We therefore calculated a 95% confidence interval, in order to obtain an error margin around the N_c_ estimate. The calculation of the 95% confidence interval of the N_c_ estimate was done using the following formulas:

$S+\frac{(Nc-S)}{C}$and $S+(Nc-S)\times C$

with:

$$C=\exp\left\{ 1.96\times\left[ \log\left( 1+\frac{\hat{var}Nc}{\left( Nc-S \right)^{2}} \right) \right]^{\frac{1}{2}} \right\}$$

and with :

$$\hat{var}Nc=F_{2}\times\left[ 0.25\times\left( \frac{F_{1}}{F_{2}} \right)^{4}+\left( \frac{F_{1}}{F_{2}} \right)^{3}+0.5\times\left( \frac{F_{1}}{F_{2}} \right)^{2} \right]$$

Thus, by considering S = 902, F1 = 782 and F2 = 120 (values extracted from the EPIDOR report; EPIDOR, Verdeyrou, Guerri, 2016), we calculated a value of N_c_ of 3,450 individuals (CI 95% = [2926.9 - 4108.3]) for the section of the Dordogne River that flows between Pessac and Mauzac.

We then estimated Ne by using a sub-sampling of individuals from our Dordogne populations that overlaps with the location where the CMR was conducted (i.e. individuals from the sampling site DOR3; Table 1 and Figure 1 in the main document). Given that we had genetic data of catfish caught in two different years in this sector (2016 and 2017; see Table 1), we used the temporal method of Jorde and Ryman (1996) implemented in the NeEstimator software (Do and al., 2014), and obtained an N_e_ of 1275.5 individuals (95% confidence interval = [841.2 - 1799]).

We finally used these N_c_ and N_e_ estimates (and their confidence intervals) to calculate an N_e_ / N_c_ ratio equal to 0.3697 (CI95% = [0.2048 – 0.615]).

**FIGURE S1:** Spatial patters of genetic variability observed at the Garonne-Dordogne basin scale for *Silurus glanis*. (A) test for the occurrence of a DIGD spatial pattern, i.e. a downstream increase in genetic diversity, measured here as the mean allelic richness (AR) across loci within sites. (B) test for the occurrence of a “mighty headwaters” spatial pattern, i.e. a pattern predicting that sites located upstream might exhibit higher genetic uniqueness (measured as Fst_UNI_). (C) test for the occurrence of an isolation-by-distance pattern. Only (C) exhibited a significant spatial pattern of genetic variation.

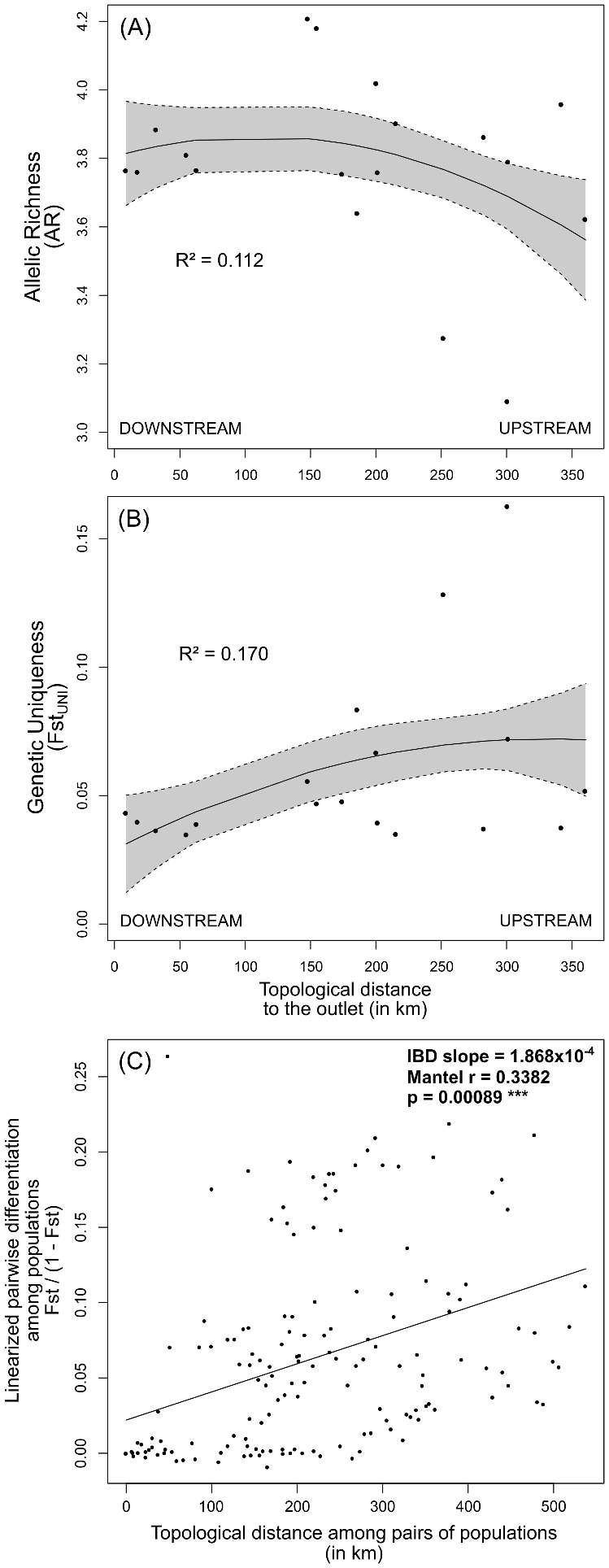

**FIGURE S2:** Cross-entropy plot to determine the best number of K clusters between 1 and 20. The vertical line shows the most probable K value, corresponding to the first elbow of the plot.

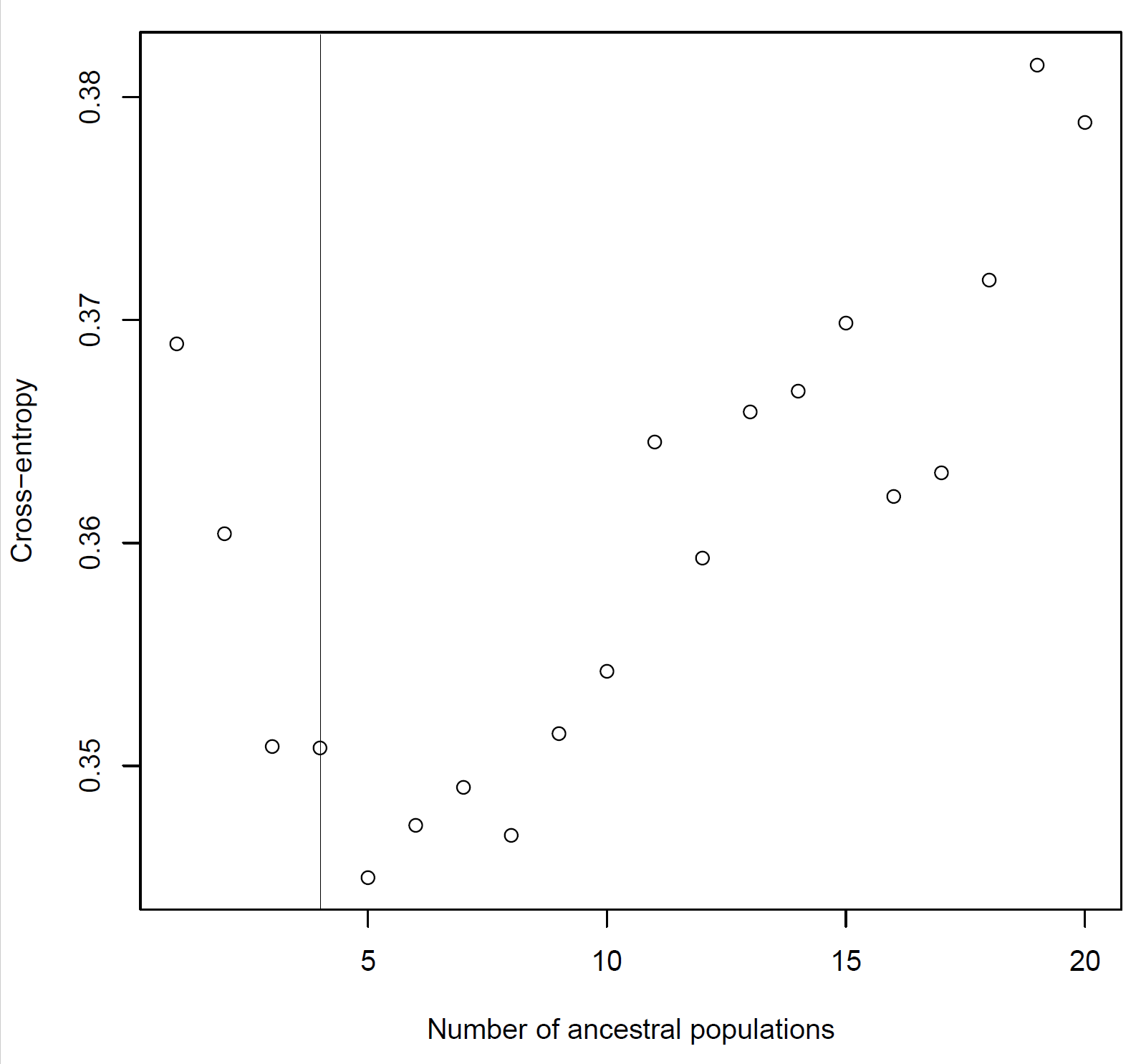
